# Structural basis of glycerophosphodiester recognition by the *Mycobacterium tuberculosis* substrate-binding protein UgpB

**DOI:** 10.1101/529602

**Authors:** Jonathan Fenn, Ridvan Nepravishta, Collette S Guy, James Harrison, Jesus Angulo, Alexander D Cameron, Elizabeth Fullam

## Abstract

*Mycobacterium tuberculosis (Mtb)* is the causative agent of tuberculosis (TB) and has evolved an incredible ability to survive latently within the human host for decades. The *Mtb* pathogen encodes for a low number of ATP-binding cassette (ABC) importers for the acquisition of carbohydrates that may reflect the nutrient poor environment within the host macrophages. *Mtb* UgpB (Rv2833) is the substrate binding domain of the UgpABCE transporter that recognises glycerophosphocholine (GPC) indicating a potential role in glycerophospholipid recycling. By using a combination of saturation transfer difference (STD) NMR and X-ray crystallography we report the structural analysis of *Mtb* UgpB complexed with GPC and have identified that *Mtb* UgpB is promiscuous for other glycerophosphodiesters. Complementary biochemical analyses and site-directed mutagenesis define the molecular basis and specificity of glycerophosphodiester recognition. Our results provide critical insights into the structural and functional role of the *Mtb* UgpB transporter and reveal that the specificity of this ABC-transporter is not limited to GPC therefore optimising the ability of *Mtb* to scavenge scarce nutrients and essential glycerophospholipid metabolites during intracellular infection.

## Introduction

Bacterial pathogens have evolved a wide range of strategies to survive and thrive within their host environment. The ability to assimilate nutrients is vital and pathogens have evolved diverse strategies to uptake and scavenge the scarce energy sources that are available to them. In the context of intracellular microbial infections there is growing evidence that in a nutrient limited environment the interplay between the host and the pathogen is important. This is manifested through the ability of bacterial pathogens to utilise discrete nutrient sources with dedicated transport machinery for import. Glycerophosphodiester metabolites that are released by the action of phospholipases on host phospholipids represent an important nutrient source for the supply of carbon and phosphate.

*Mycobacterium tuberculosis (Mtb)* is a major human pathogen and is now the leading cause of death from a single infectious agent worldwide, resulting in more deaths each year than HIV and malaria combined ^1^. *Mtb* is a highly evolved pathogen that is able to persist and survive intracellularly within macrophages for decades ^2^. However, the essential nutrients that are available to *Mtb* within the stringent environment of the human host and acquisition systems are poorly understood ^3,4^ Understanding the molecular mechanisms that enable *Mtb* to survive within this niche environment and the nutrients that are assimilated is critical to understand this major global pathogen and for the development of new therapeutic approaches.

Despite the limited availability of sugars within the macrophage environment *Mtb* is equipped with five putative importers of carbohydrate substrates: four members of the ATP-binding cassette (ABC) transporter family and one belonging to the major facilitator superfamily ^3,4^ Until recently the substrates for these transporters were unresolved, however, recent studies have demonstrated a role for the ABC-transporters in the recycling of components from the complex *Mtb* cell wall. Trehalose is recycled from the *Mtb* cell envelope glycolipid trehalose monomycolate and taken up by the LpqY-SugABC transporter, which plays a critical role in the virulence of the *Mtb* pathogen ^5^. The *Mtb* UspABC transporter has been found to recognise amino-sugars with a potential role in the uptake of *Mtb* cell-wall peptidoglycan fragments ^6^.

The role of the UgpABCE ABC-transporter is less clear, however studies of its substrate binding domain *Mtb* UgpB (Rv2833c) indicate its importance for *Mtb* survival and pathogenesis. Transposon mutagenesis studies have revealed that UgpB is essential for optimal *in vitro* growth and *in vivo Mtb* UgpB has been found to be upregulated during infection ^7,8^. *Mtb* UgpB has been shown to bind the glycerophosphocholine (GPC) head group of the membrane phospholipid phosphatidylcholine and metabolomic profiling by NMR of intact lung tissue at various stages of *Mtb* infection has revealed that the GPC metabolite increases significantly as infection progresses, with a concomitant decrease in phosphatidylcholine ^9^. However, despite the essential role of this *Mtb* transporter, the molecular mechanisms that dictate how GPC is recognised and whether other glycerophosphodiester metabolites are substrates for this ABC-transporter are currently unknown. The only crystal structure of *Mtb* UgpB is of the protein in an open conformation without substrate bound (PDB 4MFI)^10^. Some mechanistic understanding of substrate recognition can be obtained from the crystal structure of a homologue from *E. coli* with low sequence identity (25%) in complex with glycerol-3-phosphate (G3P) (PDB 4AQ4) ^11^. However*,Mtb* UgpB does not bind G3P and comparison of the closed G3P-bound *E.coli* UgpB with the open *Mtb* UgpB in the absence of substrate (PDB 4MFI) reveals notable differences in the binding sites of these homologous proteins indicating that these UgpB ABC-transporters have diverged to have different substrate specificities. This may reflect the nutritional requirements of the specific organism within different host environments and also the ability of bacteria to produce G3P extracellularly through the action of secreted glycerophosphodiesterases that hydrolyse glycerophopshodiesters ^12^. Other microorganisms that import GPC have evolved to use either permeases or proton symporters that belong to the major facilitator superfamily indicating that glycerophosphodiester uptake is not limited to ABC-transporters ^13,14^ It is likely that the divergence of transport systems for the import of glycerophosphodiesters reflects the evolutionary divergence and intracellular life-style of the pathogen and the metabolites available within its niche environment.

In this study, we report a detailed functional and structural characterisation of the *Mtb* UgpB substrate binding domain of the ABC-transporter using a combination of biochemical and biophysical approaches. We report the first crystal structure of *Mtb* UgpB in complex with GPC and identify, in both solid and solution state, the molecular determinants of binding and critical features for glycerophosphodiester recognition. Structure guided-mutagenesis has revealed the crucial role of binding-site residues that underpin substrate binding and function. Moreover, we show that *Mtb* UgpB has an unexpectedly broad selectivity for glycerophosphodiesters which highlights that the *Mtb* UgpABCE transporter uptakes metabolites derived from various glycerophospholipids. Thus, *Mtb* has evolved to use a broad spectrum of nutrients *via* a single ABC-transporter that enables it to adapt and assimilate essential nutrients during intracellular infection.

## Results

### Production of N-terminally truncated UgpB from *Mycobacterium tuberculosis*

An *N*-terminal truncated *Mtb* UgpB, corresponding to removal of residues 1-34 predicted to form a trans-membrane anchor-helix, was cloned into the pYUB1062 vector with a C-terminal hexa-histidine affinity tag and expressed in *Mycobacterium smegmatis* mc^2^4517. Soluble *Mtb* UgpB protein was obtained and purified to apparent homogeneity using Co^2+^-affinity, anion exchange and size-exclusion chromatography (Supporting information Fig. S1). The identity of the *Mtb* UgpB protein was confirmed by using in-gel trypsin digestion and analysis of the peptides by mass spectrometry.

### Co-crystal structure of *Mtb* UgpB with GPC

We crystallised the reductively methylated UgpB in the presence of GPC. The UgpB protein co-crystallised with GPC with four molecules in the asymmetric unit. Phases for the structure were determined by molecular replacement using each of the two domains from the apo-structure of *Mtb* UgpB (PDB 4MFI) as separate search models and the structure was refined at a resolution of 2.3 Å, to a *R_work_* of 20.6 % and *R_free_* of 25.6 %, see Table 1 for the data collection and refinement statistics. Structural superposition of each molecule of *Mtb* UgpB using PDBeFOLD ^15^ indicates that each molecule within the asymmetric unit is equivalent, aligning with r.m.s.d of 0.35 - 0.44 Å for 394-395 residues. The crystal packing and analysis of the packing interfaces using PDBePISA^16^ does not suggest that *Mtb* UgpB forms dimers or higher oligomers and is consistent with our analytical gel filtration studies where the protein behaves as a monomer in solution with an apparent molecular weight of 44 kDa (Supporting information, Figure S1D). It is therefore likely that the monomer is the biologically relevant unit, consistent with substrate binding domains of other ABC-transporters ^17,18^.

**Table 1.**
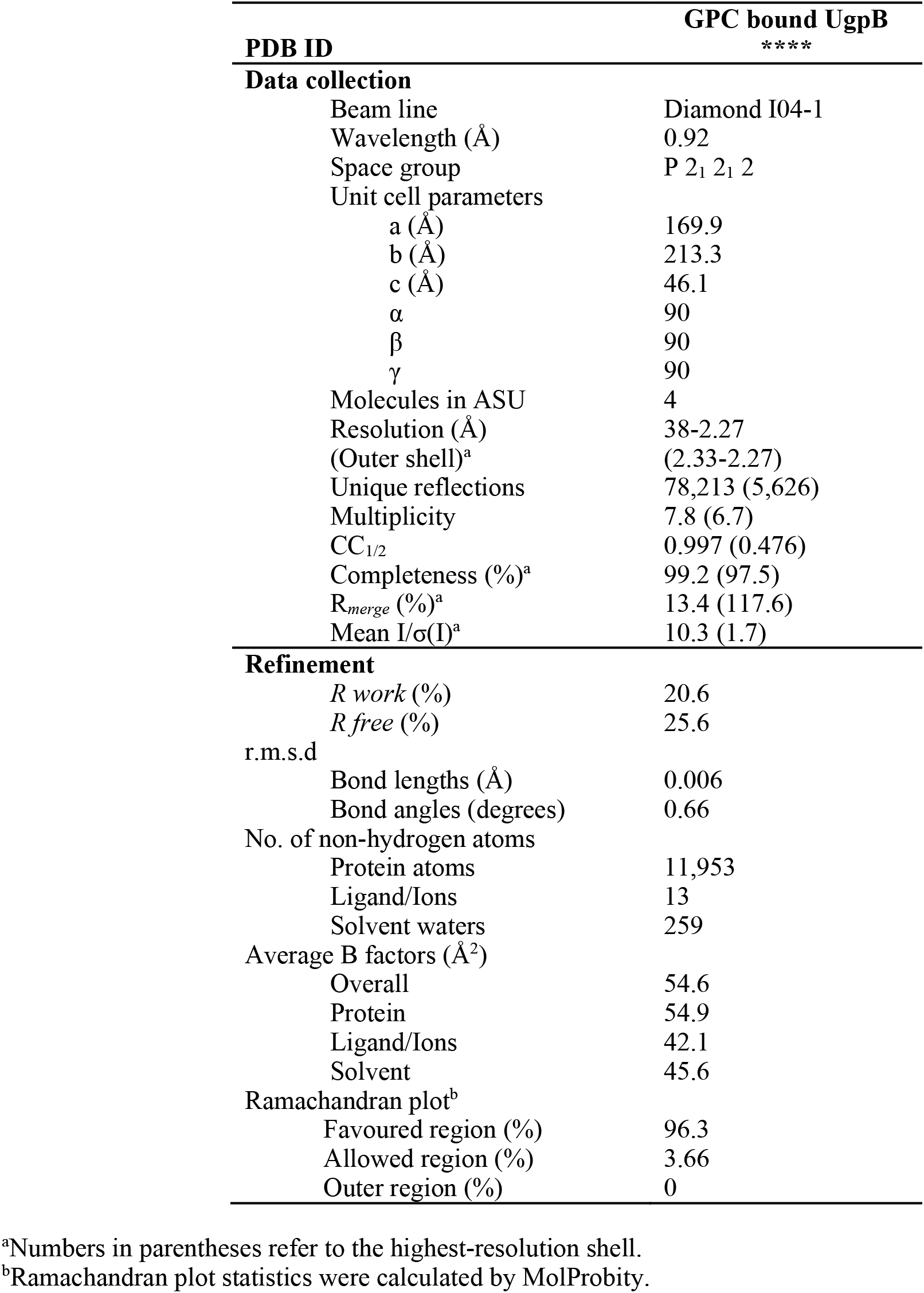
Crystallographic parameters for *Mtb* UgpB in complex with GPC

| PDB ID | GPC bound UgpB<br>**** |
| --- | --- |
| **Data collection** |  |
| Beam line | Diamond I04-1 |
| Wavelength (Å) | 0.92 |
| Space group | P 2<sub>1</sub> 2<sub>1</sub> 2 |
| Unit cell parameters |  |
| a (Å) | 169.9 |
| b (Å) | 213.3 |
| c (Å) | 46.1 |
| α | 90 |
| β | 90 |
| γ | 90 |
| Molecules in ASU | 4 |
| Resolution (Å) | 38-2.27 |
| (Outer shell)<sup>a</sup> | (2.33-2.27) |
| Unique reflections | 78,213 (5,626) |
| Multiplicity | 7.8 (6.7) |
| CC<sub>1/2</sub> | 0.997 (0.476) |
| Completeness (%)<sup>a</sup> | 99.2 (97.5) |
| R<sub>merge</sub> (%)<sup>a</sup> | 13.4 (117.6) |
| Mean I/σ(I)<sup>a</sup> | 10.3 (1.7) |
| **Refinement** |  |
| R<sub>work</sub> (%) | 20.6 |
| R<sub>free</sub> (%) | 25.6 |
| r.m.s.d |  |
| Bond lengths (Å) | 0.006 |
| Bond angles (degrees) | 0.66 |
| No. of non-hydrogen atoms |  |
| Protein atoms | 11,953 |
| Ligand/Ions | 13 |
| Solvent waters | 259 |
| Average B factors (Å<sup>2</sup>) |  |
| Overall | 54.6 |
| Protein | 54.9 |
| Ligand/Ions | 42.1 |
| Solvent | 45.6 |
| Ramachandran plot<sup>b</sup> |  |
| Favoured region (%) | 96.3 |
| Allowed region (%) | 3.66 |
| Outer region (%) | 0 |

<sup>a</sup>Numbers in parentheses refer to the highest-resolution shell.
<sup>b</sup>Ramachandran plot statistics were calculated by MolProbity.

### Overall structure of the *Mtb* UgpB-GPC complex

*Mtb* UgpB comprises two α/β domains (Figure 1). Domain I (residues 1-154 and 307-365) consists of a five-stranded β-sheet surrounded by 11 α-helices and domain II (residues 155-306 and 366-436) of a four-stranded β-sheet enclosed by 9 α-helices. The two domains, or globular lobes, are connected *via* two flexible hinges that are formed between residues Arg152-Pro155 and Ala290-Ala307. Relative to the apo crystal structure there is a 22° rotation of domain I relative to domain II about the interdomain screw-axis with three hinge/binding regions identified from DynDom analysis ^19^ (residues 152-153, 304-306 and 362-372 (Supporting information, table S1)). This bending movement results in an almost two-fold reduction in the volume of the cavity from 1986 Å^3^ to 791 Å^3^, as determined by CAVER ^20^, which is in-line with the ‘Venus Fly-trap mechanism’ for other substrate-binding proteins ^17,18^ that close when substrate is bound. Interdomain bridging and stabilisation of this closed conformation of the protein is centred around Arg385, which forms interdomain hydrogen bonds with Asp102 from domain I and Gln381 from domain II. The individual domains of *Mtb* UgpB *apo-* and GPC co-complex structures align with r.m.s.ds of 0.57 Å and 0.75 Å for domains I and II respectively (over 178 atoms, Domain I and over 216 atoms, Domain II, PDBeFOLD ^16^). In comparison, superposition of *Mtb* UgpB *apo-* and GPC co-complex structures align with a r.m.s.d. of 2.2 Å (over 385 residues) highlighting the importance of an interdomain conformational change mechanism for substrate recognition by *Mtb* UgpB.

**Figure 1.**
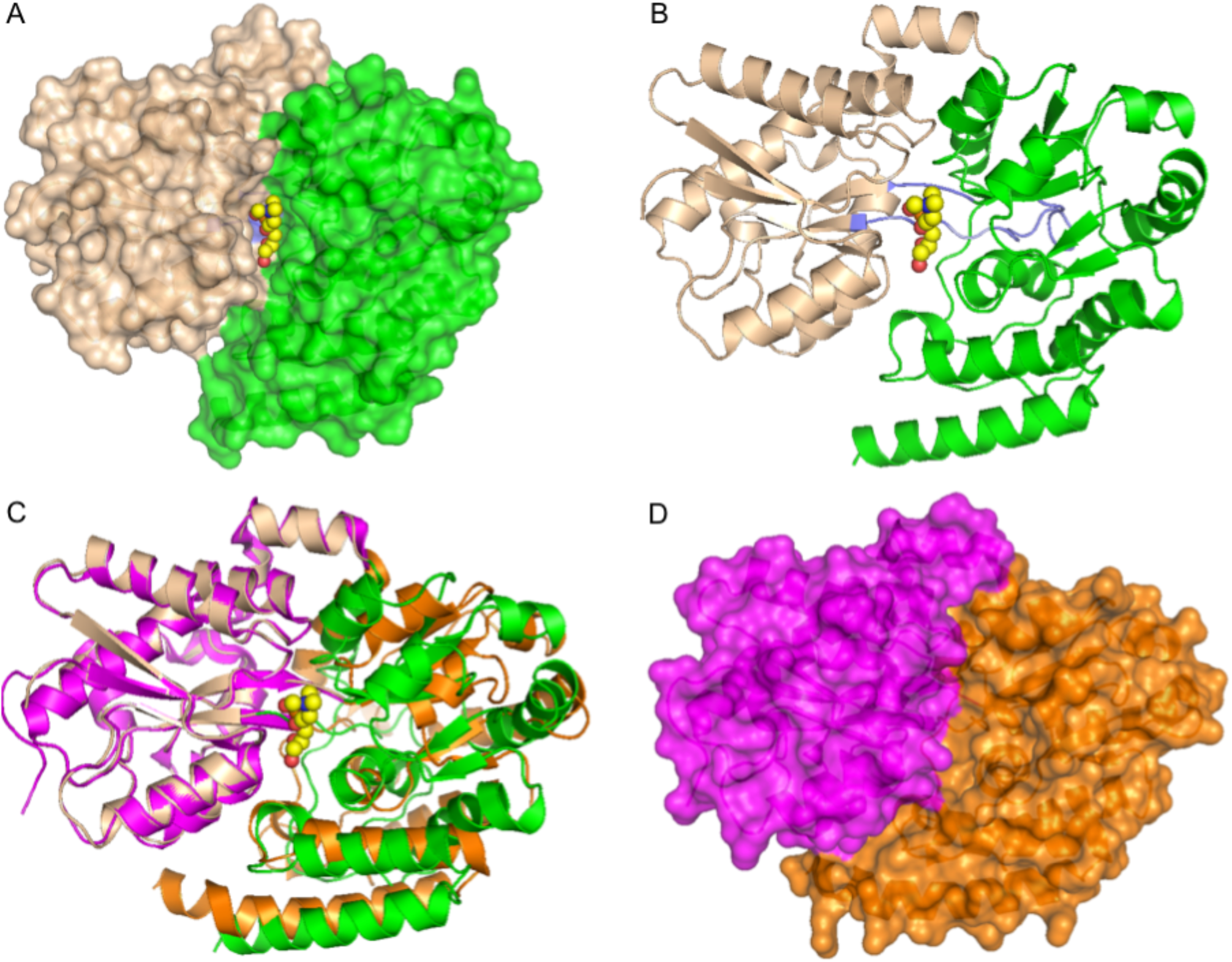
Crystal structure of *Mtb* UgpB. A) Surface representation of *Mtb* UgpB in complex with GPC. The two domains are highlighted, domain I (brown) and domain II (green). The GPC ligand is represented as spheres with yellow carbon atoms. B) Cartoon representation of *Mtb* UgpB in complex with GPC identifying the secondary structure elements. Domain I (brown), domain II (green) and the two hinge regions are highlighted in blue. The GPC ligand is represented as spheres with yellow carbon atoms C) Superposition of Domain I of GPC *Mtb* UgpB co-complex (brown/green) with Domain I of apo *Mtb* UgpB (PDB 4MFI) (magenta/orange). D) Surface representation of the unliganded *Mtb* UgpB (PDB 4MFI) with the two domains coloured magenta (Domain I) and orange (Domain II).

### The ligand-binding site of *Mtb* UgpB

Well defined electron density for the GPC ligand in all *Mtb* UgpB molecules within the crystal unit was observed enabling the GPC ligand to be modelled in the *Mtb* UgpB binding-site (Supporting information Fig. S2A). The GPC ligand is found in an identical position and orientation in each subunit (Supporting information Fig. S2B). Notably, the electrostatic surface shows that GPC is buried in the prominent, acidic interface that is formed between the two domains of UgpB and makes contact to both. The GPC is precisely orientated within the binding cleft such that the glycerol moiety is buried at the base of the cavity, in close proximity to the flexible hinge region centred around Arg385, whilst the choline moiety extends outwards towards the solvent exposed channel entrance (Figure 2).

**Figure 2.**
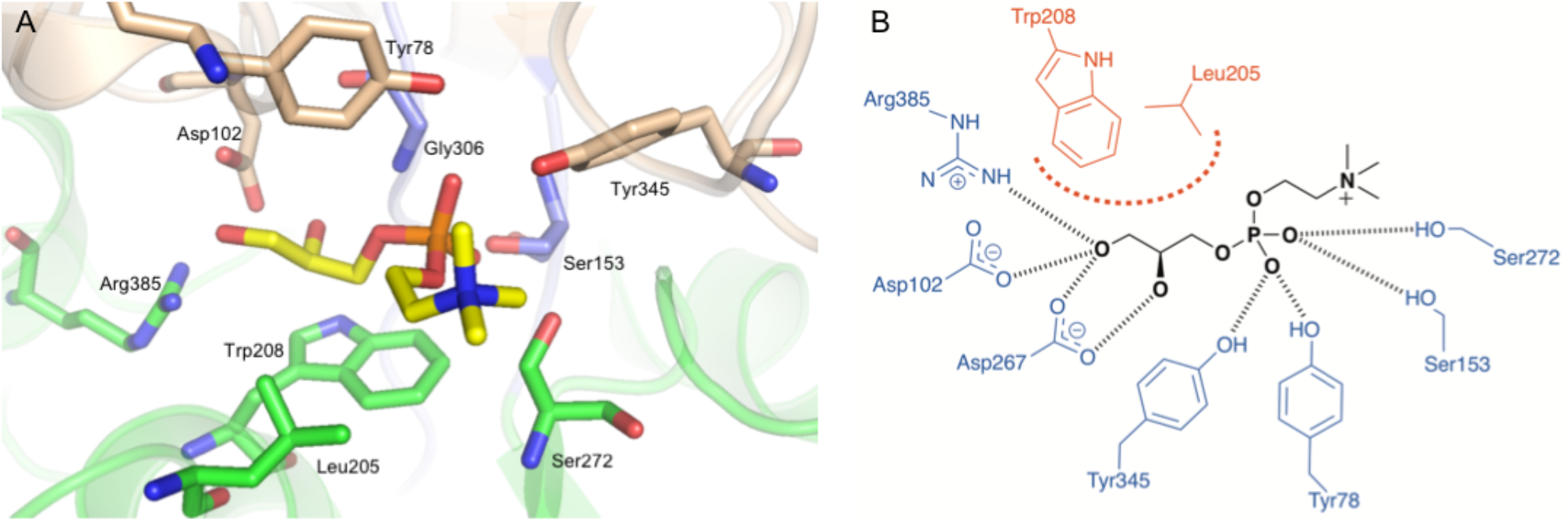
The GPC binding site in *Mtb* UgpB. A) Illustration showing GPC with yellow carbon atoms and selected *Mtb* UgpB amino acid residues in stick representation (coloured brown for residues within Domain I, and green for residues with Domain II. B) Schematic diagram of the interactions of *Mtb* UgpB with GPC. Dashed lines (black) represent hydrogen bonding, thick dotted line (red) represents hydrophobic interactions

The glycerol moiety is located between the side chains of Leu205 and Trp208 from domain II (Figure 2). The ring system of Trp208 lies approximately parallel to the C1, C2 and 2-hydroxy group of the glycerol moiety enabling π-stacking interactions, whilst Leu205 is orientated perpendicular to this plane and provides additional stabilisation. There is an important network of hydrogen bonding interactions that anchors GPC in the binding-pocket. The side chain of Asp102, from domain I, is orientated to enable direct hydrogen bonding to both the 1- and 2-hydroxy groups of the glycerol moiety. Two residues that comprise the flexible hinge-linkages are able to directly interact with GPC through the formation of additional hydrogen bond interactions between the side chain of Arg385 and the 1-hydroxy group and the backbone amide nitrogen atom of Gly306 with the 2-hydroxy group respectively. The direct interaction of these flexible-hinge linkages with the GPC ligand may help to stabilise the UgpB-GPC complex in the closed conformation. The phosphate group of GPC is stabilised through hydrogen bond interactions with the side chains of Tyr78 and Tyr345 (domain I), Ser153 (domain I), Ser272 (domain II) and the backbone amide of Gly306. It is striking that there are no direct or charged interactions between *Mtb* UgpB and the positively charged choline moiety, though this moiety is well-defined in the electron density.

### Comparison with the binding site of *E. coli* UgpB

Comparison with UgpB from *E coli* ^11^ indicates that the overall architecture of these two periplasmic binding proteins in complex with substrate is similar, with a r.m.s.d. of 2.1 Å (PDBeFold ^15^, target residues: 394, sequence identity 25 % (Supporting information Fig. S3), PDB code 4AQ4), Fig 3. Whilst *Mtb* was crystallised with GPC, the *E coli* protein was crystallised with G3P that we, as well as previous studies ^11^, show does not bind to *Mtb* UgpB. It is interesting to note that in both structures the common G3P core of the ligands align almost identically (Fig. 3B). However, whilst the substrate binding pocket of *Mtb* UpgB resembles that of *E. coli* UgpB there are several important differences. Notably, there are substitutions of critical residues involved in substrate binding. Leu205 is specific to *Mtb* and is replaced by a larger indole-side chain from a tryptophan residue (Trp169) in *E. coli* UgpB. In addition, *Mtb* UgpB Asp102 is replaced in *E. coli* UgpB by a glutamic acid residue (Glu66) (Fig. 3C). In this instance, the difference in the length of these acidic side-chains may influence substrate selectivity between the different organisms. Intriguingly, whilst the interaction with an arginine residue is conserved between *Mtb* and *E. coli* the arginine residues in the two proteins originate from different regions of the protein indicating an evolutionary divergence of these substrate-binding proteins. In addition, a narrowing of the *E. coli* UgpB binding cleft results from two different loop regions. One loop region (Gly221-Asp230) in domain II of *E. coli* UgpB linking α-helices 10 and 11 narrows the substrate binding cavity as a result of a 5 Å translational shift. The difference in position of a second loop comprised of residues His8-Gly12 results in the translation of the first α-helix of *E. coli* UgpB (residues 12-30) located in domain I by approximately 6 Å towards α-helix 11 of domain II which further narrows the *E. coli* UgpB substrate binding channel (Fig. 3D/E). Comparison of the region at the entrance to the binding cleft reveals an expanded pocket for *Mtb* UgpB. It is of interest to note that in chain B of *Mtb* UgpB we observe an additional glycerol molecule located in this expanded pocket that is within 4 Å of the choline moiety of GPC (Supporting information Fig. S4). A glycerol molecule is also present in the *E. coli* UgpB-G3P complex, though at a different position, indicating that for both proteins the binding pockets are larger than the recognised GPC substrate ^11^. This may be functionally significant in substrate recognition and have an important role in the accommodation and binding of alternative phosphodiester substrates.

**Figure 3.**
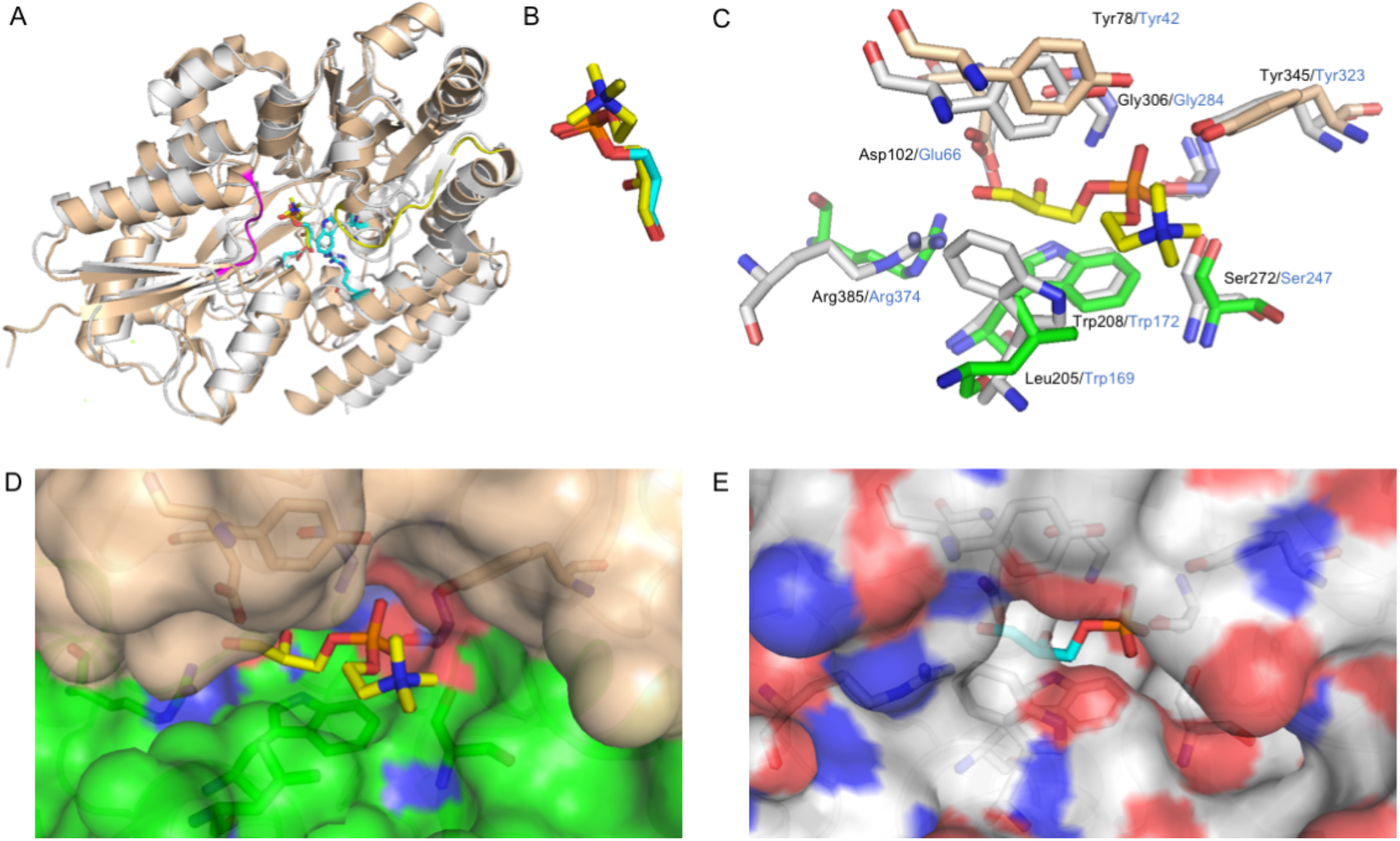
Comparison of *Mtb* UgpB with *E. coli* UgpB. A) Superposition of the *Mtb* UgpB GPC complex structure (brown) with *E. coli* UgpB in complex with G3P (PDB 4AQ4) (grey). Loop regions that differ are highlighted in yellow and magenta. B) Close-up illustration showing the binding orientation of the GPC ligand and G3P ligand in stick representation (yellow carbon atoms, GPC, cyan carbon atoms G3P) C) Close-up of the overlay of the binding-sites of GPC (*Mtb*) and G3P (*E. coli*). Selected residues are shown as sticks (*Mtb* brown, *E. coli* grey) and the font labelled in black (*Mtb*) and blue (*E. coli*). D) Surface representation of the *Mtb* UgpB GPC binding pocket with the GPC ligand in stick representation. E) Surface representation of the *E. coli* UgpB G3P binding pocket in the same orientation as D with the G3P ligand in stick representation.

### Solution saturation transfer difference (STD) NMR of *Mtb* UgpB with glycerophosphocholine

Given the apparent discrepancy between the lack of interactions formed between the choline moiety and its importance in binding, given that G3P lacking the choline moiety does not bind, we decided to investigate binding in the solution state. We employed saturation transfer difference (STD) NMR to obtain quantitative maps of the ligand-protein complex in solution (Fig. 4) ^21^. Binding was detected for GPC and binding epitope mapping was obtained and analysed as described in the methods section ^22^. The STD NMR signals and the GPC binding epitope and maps obtained are shown in Fig. 4. From the epitope map, the glycerol moiety of GPC is identified as the main recognition element showing the highest STD normalized values. In particular, the highest STD intensity values were observed for the protons in position 1 and 2 (H1G and H2G) of the glycerol moiety (Fig. 4B), with slightly lower intensity values for the protons in position 3 (H3G). The STD values decrease from the glycerol moiety to the choline group, indicating that the ligand-protein contacts are closer with the glycerol group than with choline. Intermediate and low STD NMR intensity values were observed for the protons in position 1 and 2 (H1C and H2C) while low intensity values were observed for the methyl groups from the choline moiety. A quantitative comparison of the NMR solution data with the X-ray structure of the complex was carried out using CORCEMA-ST calculations ^23^, as well as the newly developed method DEEP-STD NMR ^24^ and the results are summarised in Fig. 4. An NOE R-factor ^25^ of 0.25 was obtained when comparing the CORCEMA-ST calculated STD NMR intensities using the crystal structure with the experimentally obtained solution data. This indicates a very good agreement of the complex in solution state with the crystal structure. In order to probe for additional structural information in the solution state we then utilised differential epitope mapping by STD NMR (DEEP-STD NMR). This methodology allows us to gain information about the orientation of the ligand within the architecture of the binding site and indirectly gives information about the type of amino acids (aromatic, polar or apolar residues) surrounding the ligand in the bound state ^26^. The DEEP-STD NMR factors clearly identified that the protons in position 3 of the glycerol moiety of GPC are orientated towards aliphatic amino acids whilst the protons in position 1 in the choline moiety are oriented toward aromatic residues (Fig. 4D). Based on the crystal structure of *Mtb* UgpB these residues can be mapped to Leu205, Tyr78 and Tyr345 respectively (Fig. 2). Notably, our data shows strong correlation for the molecular determinants of GPC ligand binding to *Mtb* UgpB to GPC in both solution and solid state.

**Figure 4.**
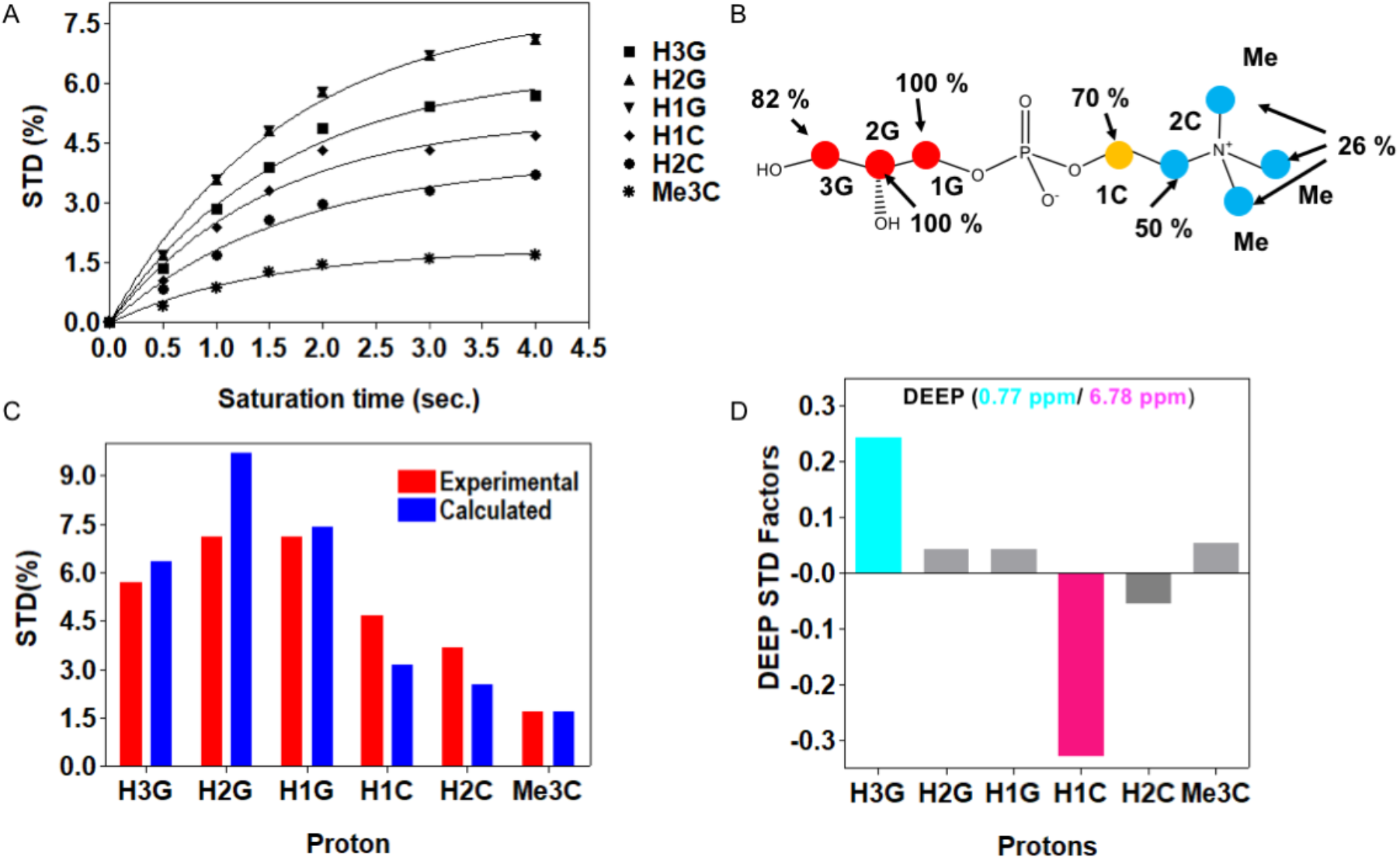
STD-NMR for *Mtb* with GPC. A) Experimental STD build up curves for the *Mtb* UgpB/GPC complex obtained as described in material and methods. B) Obtained epitope map of GPC/Mtb UgpB interaction in solution state. C) STD in red bars obtained with a 4s saturation time while in blue bars the CORCEMA-ST calculated STD from the 3D crystallographic structure of the *Mtb* UgpB/GPC complex obtained for the same saturation time. RNOE factor 0.25. D) DEEP-STD factors showing the orientation of the protons for the GPC ligand.

### Substrate specificity of *Mtb* UgpB

To establish the importance of both the polar head group and the glycerol moiety for substrate recognition binding we analysed the binding interactions of *Mtb* UgpB with G3P, the preferred substrate of *E. coli* UgpB, and phosphocholine. In contrast to GPC, no binding interactions were observed for these smaller derivatives. Taken together with our structural studies, these results indicate that whilst the glycerol moiety is the main recognition element for *Mtb* UgpB and that there are minimal interactions with the polar head group, the entire phosphodiester moiety is critical for substrate recognition and binding.

Our structural studies in both the solid and solution state revealed that the GPC substrate interacts predominantly with *Mtb* UgpB through interactions with the glycerol backbone. The lack of specific interactions between the protein and the polar choline head group located at the entrance of the substrate binding pocket led us to speculate that *Mtb* UpgB may recognise alternative glycerophosphodiester analogues. To directly investigate the substrate specificity of *Mtb* UgpB we used microscale thermophoresis (MST) to analyse the binding interactions of other phosphodiester products formed from the lipolysis of membrane glycerophospholipids (Fig. 5). From the substrates tested, in each case we were able to detect binding for GPC, glycerophosphoserine (GPS), glycerophosphoethanolamine (GPE), glycerophosphoinositol (GPI) and glycoerphosphoinositol-4-phosphate (GPI4P), (Table 2, Fig. 6). The measured *K_d_* value for GPC was consistent with previous results obtained by isothermal titration calorimetry (ITC) ^10^. Notably, *Mtb* UgpB also binds and recognises GPE, GPS, GPI and GPI4P glycerophosphodiesters with binding affinities in the micromolar range (Table 2) with a preference for positively charged polar head groups. Together, this suggests that *Mtb* has evolved to have a single ABC-transporter to scavenge a range of glycerophosphodiesters within its nutrient poor intracellular environment.

**Figure 5.**
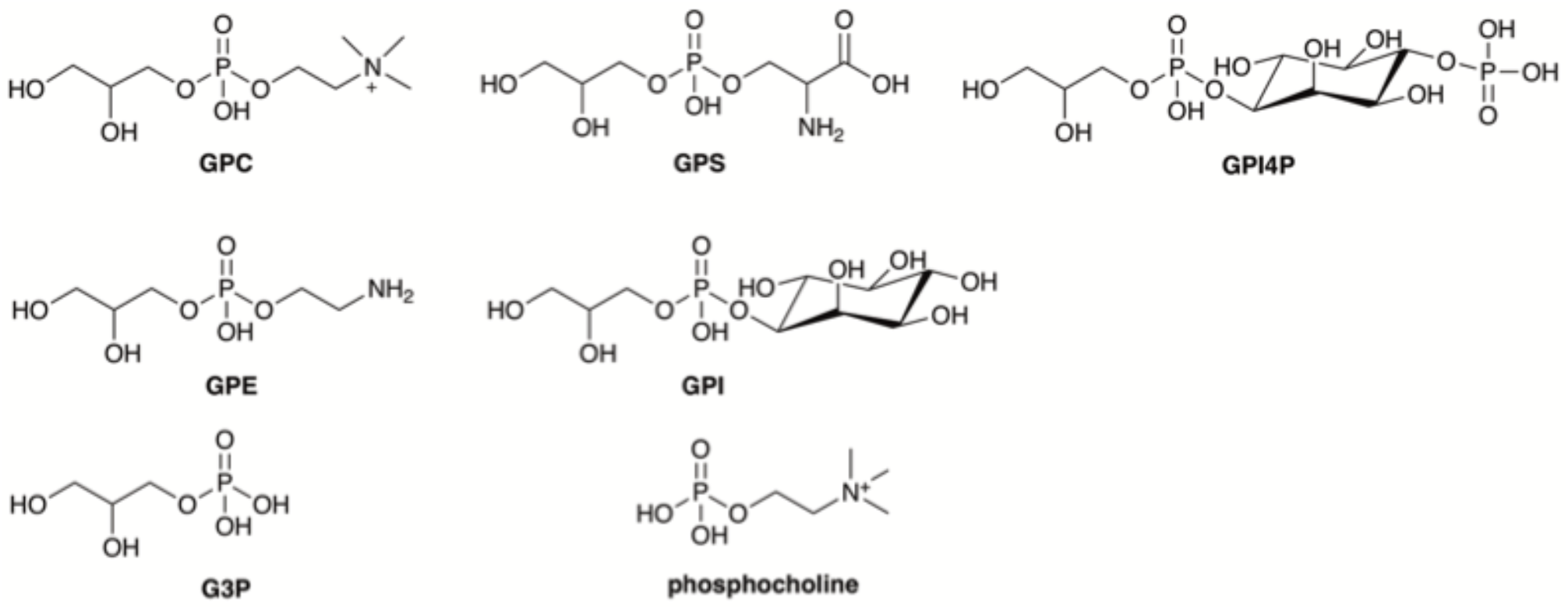
Structure of glycerophosphodiesters and derivatives probed in this study.

**Figure 6.**
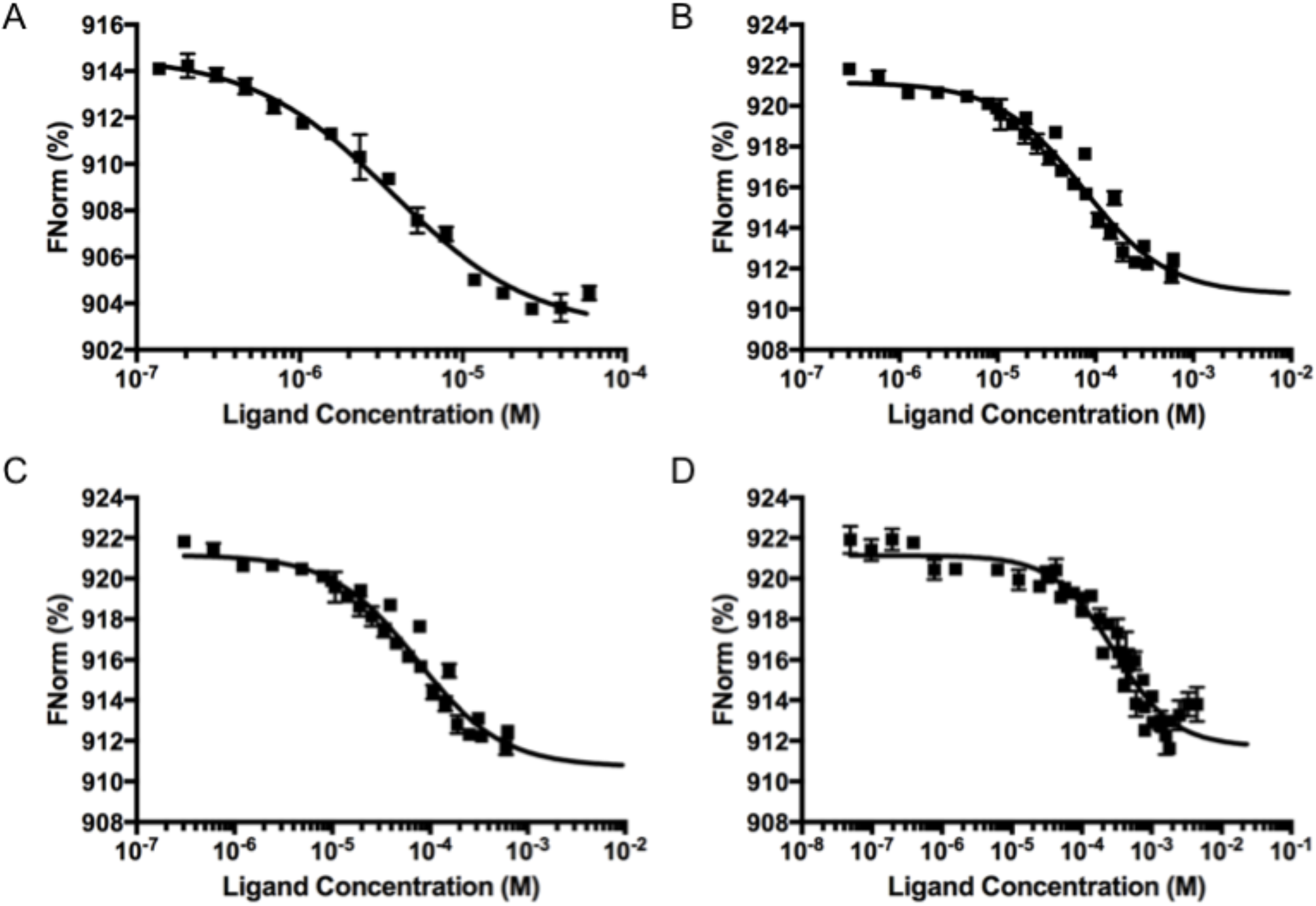
Binding affinities for *Mtb* UgpB. Binding of A) GPC, B) GPS, C) GPE and D) GPI4P to *Mtb* UgpB measured by microscale thermophoresis (MST). Error bars represent standard deviations from at least three independent experiments.

**Table 2.** Binding data for *Mtb* UgpB

| Enzyme | Substrate | $K_d(\mu\text{M})$ | Reference |
| --- | --- | --- | --- |
| *Mtb* UgpB | GPC | $3.6 \pm 0.5$ | This study |
| *Mtb* UgpB | GPS | $14.9 \pm 1.6$ | This study |
| *Mtb* UgpB | GPE | $74.7 \pm 13.9$ | This study |
| *Mtb* UgpB | GPI | $1053.2 \pm 313.4$ | This study |
| *Mtb* UgpB | GPI4P | $289.8 \pm 54.1$ | This study |
| *Mtb* UgpB | G3P | - | This study |
| *Mtb* UgpB | phosphocholine | - | This study |
| *Mtb* UgpB Tyr78Ala | GPC | - | This study |
| *Mtb* UgpB Tyr78Ala | GPS | - | This study |
| *Mtb* UgpB Tyr78Ala | GPE | - | This study |
| *Mtb* UgpB Asp102Ala | GPC | - | This study |
| *Mtb* UgpB Asp102Ala | GPS | - | This study |
| *Mtb* UgpB Asp102Ala | GPE | - | This study |
| *Mtb* UgpB Ser153Ala | GPC | $309.8 \pm 56.1$ | This study |
| *Mtb* UgpB Ser153Ala | GPS | $102.5 \pm 16.4$ | This study |
| *Mtb* UgpB Ser153Ala | GPE | - | This study |
| *Mtb* UgpB Leu205Ala | GPC | $161.7 \pm 15.9$ | This study |
| *Mtb* UgpB Leu205Ala | GPE | $1360 \pm 210$ | |
| *Mtb* UgpB Trp208Ala | GPC | - | This study |
| *Mtb* UgpB Ser272Ala | GPC | - | This study |
| *Mtb* UgpB Tyr345Ala | GPC | - | This study |
| *Mtb* UgpB Arg385Ala | GPC | - | This study |
| *Mtb* UgpB | GPC | $27.3 \pm 2.0$ | 10 |
| *Mtb* UgpB | G3P | - | 10 |
| *Mtb* UgpB | Maltose | - | 10 |
| *Mtb* UgpB Leu205Trp | GPC | - | 10 |
| *Mtb* UgpB Leu205Trp | G3P | - | 10 |
| *E. coli* UgpB | GPC | $5.1 \pm 0.3$ | 11 |
| *E. coli* UgpB | G3P | $0.68 \pm 0.02$ | 11 |

(-) = no binding detected, standard deviations from at least three independent experiments GPC: glycerophosphocholine, GPS, glycerophosphoserine, GPE: glycerophosphoethanolamine, GPI Glycerophosphoinositol, GPI4P:glycerophosphoinositol-4-phosphate.

As a final evaluation for potential substrate promiscuity we screened a panel of carbohydrates and amino acids using a thermal shift assay and assessed the binding of putative ligands that resulted in a change in the melting temperature *(T_m_)* of *Mtb* UgpB which can be indicative of binding. In total 37 potential substrates were probed, including trehalose which is known to be a substrate of the *Mtb* LpqY-SugABC ABC-transporter ^5^, and we found that none of the ligands that were screened significantly influenced the melting temperature (Supporting information, Fig S5). It appears that although *Mtb* encodes for only five putative carbohydrate importers, each transport system has a defined substrate preference. Taken together, these data indicate that the substrate binding-pocket of *Mtb* UgpB can efficiently accommodate glycerophosphodiesters but that it is not able to recognise other carbohydrates or amino acids.

### STD NMR of *Mtb* UgpB with GPI4P

Next, to validate some of the MST-binding data we used STD NMR spectroscopy for a more in-depth investigation of GPI4P binding to *Mtb* UgpB. Again the glycerol moiety of GPI4P was the main recognition element with close contacts to *Mtb* UgpB. High STD NMR intensity values were also observed for the H1 and H2 protons of the inositol ring with intermediate STD-NMR values for H3 and H4 protons and low values for H5 and H6 protons (Fig 7A, B). This differs from the situation of the choline head group of GPC where instead low STD intensities were observed. Furthermore, the DEEP-STD NMR maps reveal a slight modification in the binding orientation of the glycerol moiety of GPI4P compared to GPC as protons in position 3 orientated towards aromatic residues this time. To gain 3D structural insights about this interaction we carried out docking calculations using Autodock Vina ^27^ followed by validation using CORCEMA-ST calculations (Fig 7C). An NOE R-factor of 0.31 was obtained by comparing the CORCEMA-ST calculated STD intensities from the best scored docked structure of GPIP4 bound to *Mtb* UgpB and the experimental STD values. This indicates a good agreement of the proposed docking structure of the *Mtb* UgpB/GPIP4 complex with the experimental STD NMR data. From Fig. 7 we can observe that the protons in position 3 (H3G) are oriented toward the aromatic residues, which was also determined from DEEP-STD factors analysis. Further, also the protons of inositol-phosphate moiety are in line with the observed orientation from DEEP-STD factor analysis. In fact protons H4I, H1G, H2G are oriented toward aliphatic residue Leu205, while protons H1I, H3G, H6I, H5I are oriented toward the aromatic residues Tyr78 and Tyr345, validating the proposed model structure with the experimental STD and DEEP-STD NMR data.

**Figure 7.**
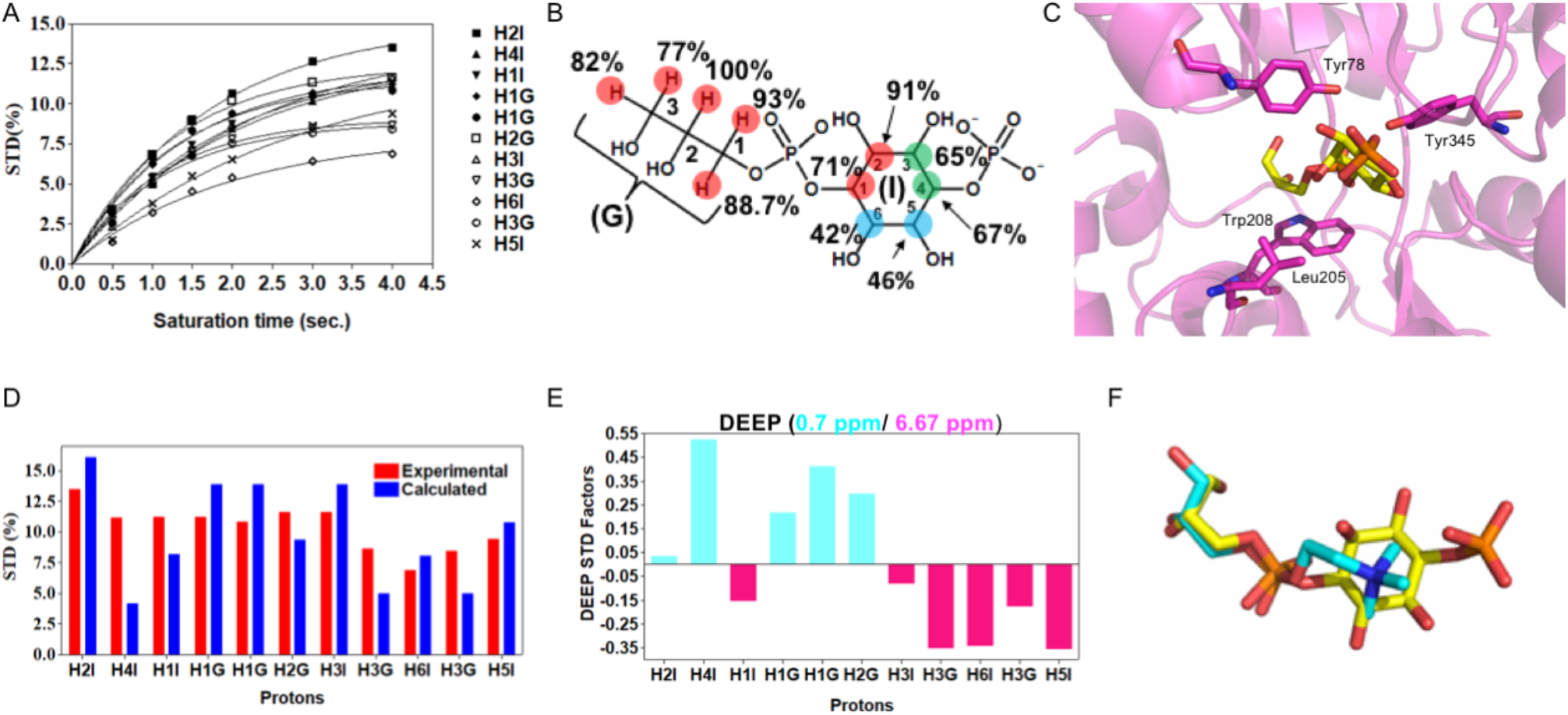
STD NMR of *Mtb* UgpB with GPI4P. **A)** STD build up curve for the *Mtb* UgpB/GPIP4 complex obtained as described in material and methods. **B)** Obtained epitope map of GPIP4/Mtb UgpB interaction in solution state. **C)** Docked structure of the GPIP4 in the binding site of *Mtb* UgpB. GPI4P is in stick representation with the carbon atoms in yellow. The binding orientation of GPC obtained from the crystal structure is shown in stick representation with orange carbon atoms. D) Experimental STD in red bars obtained with a 4s saturation time while in blue bars the CORCEMA-ST calculated STD from the 3D docked structure of the *Mtb* UgpB/GPIP4 complex obtained for the same saturation time. RNOE factor 0.31 E) DEEP-STD factors showing the orientation of the protons of GPIP4. **F)** Close-up overlay of the binding orientations of GPC (cyan carbon atoms) with GPI4P (yellow carbon atoms).

### Activity of sequence variants

In order to complement our structural studies in both the solution and solid state and assess the significance of individual amino acids that were identified to be important in molecular recognition and binding we introduced single point mutations in eight individual residues that were suggested to interact with the glycerophosphodiester ligands. In each case, we confirmed that the substituted alanine mutation was not detrimental to the correct folding of the protein by circular dichroism spectroscopy (Supporting Information Fig. S6). MST was used to determine the binding affinities of the *Mtb* UpgB protein with GPC and complete abrogation of binding was observed when Tyr78, Asp102, Trp208, Ser272, Tyr345 and Arg385 were individually replaced by an alanine, confirming the significance of these residues in substrate selectivity and importance in binding recognition. In contrast, binding of GPC was still observed when Ser153 and Leu205 were replaced by alanine, with a corresponding 85- and 45-fold reduction in the *K_d_* values respectively (Table 2), indicating that whilst these two individual residues are important for binding, they are not critical. Failure of these single-residue mutants to completely abolish binding reflects that multiple amino-acids are involved in the interaction with GPC, as observed from the crystal structure. Previous studies that mutated *Mtb* UgpB Leu205 to a tryptophan residue to mimic the situation found in *E. coli* UgpB were detrimental for binding of GPC, indicating that the bulky indole side-chain cannot be tolerated in *Mtb* UgpB ^10^ and did not enable recognition of G3P. The distinct glycerophosphodiester-recognition of *Mtb* UgpB compared with *E.coli* UgpB indicates that the mycobacterial UgpB transporter has evolved to have unique specificity and function that is distinct from other UgpB proteins.

## Discussion

The ongoing battle of *Mtb* to assimilate scarce nutrients during intracellular infection is a critical factor for the survival of this major global pathogen. Thus, the discovery that the *Mtb* UgpABCE ABC-transporter binds and recognises a range of glycerophosphodiesters as a route to obtaining scarce resources within its niche host environment is a significant step forward to understanding the mechanisms of nutrient import and the energy sources that are available.

Altogether, our biochemical, X-ray crystallographic and STD-NMR data are consistent and provide the first direct evidence that *Mtb* UgpB recognises and binds a broad spectrum of glycerophosphodiesters. This is significant and provides new insights into the substrate repertoire of this essential *Mtb* ABC-transporter. Previous studies have found that the *Mtb* UgpB substrate binding domain has specificity for GPC whilst the potential to recognise other physiological ligands was not explored ^10^. Our biochemical analysis clearly reveals that *Mtb* UgpB can specifically recognise three additional glycerophosphodiesters: GPS, GPE and GPI4P. Although there is a clear preference and tighter binding for the choline glycerophosphodiester derivative, the serine and ethanolamine head groups are also well tolerated. Replacement of the polar head group with inositol-4-phosphate, in the case of GPI4P, resulted in a reduction in the observed binding affinities, although it is still in the micromolar range and comparable with reported affinities for other carbohydrate-binding proteins of ABC-transporters ^18^. In contrast the inositol analogue that lacks the phosphate group is only moderately tolerated. In addition, both the polar head group and glycerol moieties are required for substrate recognition demonstrated by a loss of binding for G3P or phosphocholine, indicating that *Mtb* UgpB does not have the capacity to recognise smaller glycerophosphodiester metabolites The preference for GPC could suggest that as phosphatidylcholine is the main glycerophospholipid in human lung tissue ^28^ that *Mtb* UgpB has evolved to recognise the most abundant glycerophosphodiester available within the host environment, with the potential to recognise and transport a spectrum of additional glycerophosphodiesters depending on the growth conditions and nutrient availability during intracellular infection. Notably, from the extensive panel of carbohydrates and amino acids also tested in the study we identified that *Mtb* UgpB does not have the capacity to recognise any additional substrates, highlighting its unique specificity for glycerophosphodiesters and reinforcing the notion that the *Mtb* transporters have distinct substrate preferences ^3,5,6^.

Importantly, in these studies we were able to determine molecular-level insights into the binding specificity of *Mtb* UgpB for glycerophosphodiesters. To our knowledge, this constitutes the first structure of a substrate-binding protein bound to GPC. Specifically, we were able to unambiguously determine the position and orientation of the GPC and GPI4P substrates in both solid- and solution-state through a combination of X-ray crystallography and STD-NMR. The binding mode of the G3P core of GPC resembles the situation found in *E. coli* UgpB in complex with G3P ^11^, even though *Mtb* UgpB is unable to bind or recognise this smaller G3P ligand. The substrate preference of *Mtb* UgpB is manifested through a distinct binding pocket that has a pronounced effect on substrate recognition, with all structure-guided point-mutations within the binding-cleft resulting in either a marked reduction or complete loss of binding. Remarkably, we show that the glycerol moiety is the main recognition element and that there are minimal interactions with the polar head group.

Despite the lack of direct contacts between the polar head group of the glycerophosphodiester and *Mtb* UgpB there is a specific requirement for a glycerophosphodiester ligand and this defines substrate binding and recognition. The critical influence of the head group on substrate selectivity supports the conclusion that the size and charge of this moiety affects the mode of binding that is altered through a re-orientation of the glycerol tail.

It is particularly interesting to highlight the inability of *Mtb* UgpB to bind G3P as this feature differs significantly from *E. coli* UgpB, which binds G3P and GPC with comparable affinities ^11^. Consistent with the low sequence identity between these substrate-binding domains (25%), there are significant differences between the binding pockets of these two organisms and propose that *Mtb* has evolved unique structural features to facilitate the import of glycerophosphodiesters present in its niche host environment. The lack of recognition of G3P by *Mtb* UgpB is consistent with the intracellular location of two putative *Mtb* glycerophosphodiesterase enzymes (GlpQ: GlpQ1, Rv3842c; GlpQ2, Rv03127c) that are predicted to degrade glyercophosphodiesters to produce G3P and the corresponding alcohol ^29,30^. In direct contrast *E. coli*, and other bacteria, including *Staphylococcus aureus*, secrete glycerophosphodiesterase enzymes to enable the extracellular production of G3P and this is consistent with the ability of the periplasmic *E. coli* UgpB to recognise the G3P metabolite ^12,31,32^.

The recycling and uptake of essential metabolites is emerging as an essential feature of *Mtb* survival during intracellular infection. It is likely that glycerophosphodiesters imported by the *Mtb* UgpABCE ABC-transporter are derived from the degradation of host lipids as a major sources of carbon and phosphate and that either the host or *Mtb* phospholipases act in concert to generate these essential energy sources ^29,33–36^. Notably, these glycerophospholipids are also major constituents of the *Mtb* cell envelope ^37,38^. The *Mtb* inner-membrane is comprised of the glycerophospholipids: phosphatidylinositol (PI) and mannosylated phosphatidylinositol, phosphatidylglycerol, phosphatidylserine and phosphatidylethanolamine. In addition, mannosylated phosphatidylinositol and phosphatidylethanolamine are constituents of the outer layer of the *Mtb* cell envelope. Given this scenario, it is possible that glycerophosphodiester metabolites are also derived through remodelling of *Mtb* cell envelope ^33,34,36,39,40^. Further experiments are underway to elucidate the involvement of host and Mtb-derived lipids.

In conclusion, to date, the nutrient requirements of *Mtb* during infection and the corresponding transport systems have not been fully elucidated. The structural and functional understanding of mycobacterial ABC-transporters that import essential nutrients is an important step to understanding the mechanisms that support intracellular survival. Importantly, we have identified that the essential *Mtb* UgpABCE importer is linked with glycerophosphodiester uptake with wide substrate selectivity. For the first time, we have established the molecular determinants of the distinct substrate selectivity of the UgpB substrate binding protein from the *Mtb* pathogen that has important structural and functional differences with *E. coli* UgpB. We therefore propose a new role for the *Mtb* UgpABCE transporter in the uptake of glycerophosphodiesters generated from the degradation of membrane phospholipids as a route to scavenge scarce nutrients during intracellular infection.

## Methods

### Materials and reagents

All chemicals and reagents were purchased from Sigma-Aldrich, unless specified. PCR and restriction enzymes were obtained from New England Biolabs. Double-distilled water was used throughout.

### Plasmid construction

An *N*-terminal truncated form (codons 35-436) of the *Mtb ugpB* gene (Rv2833c) was amplified from *Mtb* genomic DNA by PCR using gene specific primers listed in Supporting Table S2. The PCR amplification (Q5 polymerase (NEB)) consisted of 30 cycles (95°C, 2 min; 95°C, 1 min; 60°C, 30 s; 72°C, 3 min), followed by an extension cycle (10 min at 72°C). The resulting PCR product was cloned into the vector pYUB1062 using the *NdeI* and *HindIII* restriction enzyme sites resulting in the construct *ugpB-pYUB1062*. Targeted single-site substitutions were introduced into *ugpB-pYUB1062* using the primers that are detailed in Supporting Table S2, with Phusion HF polymerase and the PCR cycle (98°C, 30 s; 20 cycles of 98°C, 30 s; 60°C, 30 s; 72°C, 4 min; followed by 5 min at 72°C), followed by digestion with 1μL DpnI. All plasmid sequences were verified by DNA sequencing (GATC) and used for protein expression.

### Heterologous overexpression of *Mtb* UgpB

*Mycobacterium smegmatis* mc^2^4517 competent cells were transformed with the appropriate *ugpB-pYUB1062* expression plasmid and grown at 37 °C to an optical density at 600 nm (OD600) of 0.4 in LB media supplemented with 0.05% Tween-80, 0.2% glycerol, 100μg/mL hygromycin and 25μg/mL kanamycin. Protein production was induced with 0.2% acetamide and the culture was grown at 37 °C for 20 hours with shaking at 180 rpm. The cells were harvested and resuspended in lysis buffer (25 mM NaH2PO4, 500 mM NaCl pH 7.4 (Buffer A), supplemented with 0.1% Triton-X 100, DNAse and Complete Protease Inhibitor Cocktail (Pierce)). The cells were freeze-thawed and sonicated on ice. Following centrifugation (39,200 g, 40 min, 4°C) the supernatant was loaded onto a pre-equilibrated Co^2+^-affinity resin (HisPure). The *Mtb* UgpB protein was eluted from the Co^2+^-affinity column in buffer A with increasing concentrations of imidazole. Fractions containing the protein, as determined by SDS-PAGE, were pooled and dialysed against buffer B (25 mM HEPES, 150 mM NaCl, pH 7.0) at 4 °C for 16 hours. Following dialysis, the protein was loaded onto a pre-equilibrated QHP ion exchange column (GE Healthcare) and eluted with buffer B containing increasing concentrations of NaCl (0.1-1 M). Fractions containing *Mtb* UgpB were pooled and loaded onto a Supderdex 75 pg HiLoad 16/600 gel filtration column (GE Healthcare) and eluted with buffer C (25 mM HEPES, 150 mM NaCl, 10% glycerol pH 7.0). The fractions that contained *Mtb* UgpB were combined and concentrated by ultrafiltration (10 kDa cut-off, Amicon Ultra) to ∼ 5-10 mg/mL prior to storage at −80°C. The identity of the protein was confirmed by tryptic digest and nanoLC-ESI-MS/MS (WPH Proteomics facility, University of Warwick).

### Circular Dichroism (CD) analysis

Purified *Mtb* UgpB proteins were diluted to 0.25 mg/mL and dialysed in the following buffer: 25 mM NaH2PO4, 100mM NaCl, 10% glycerol pH 7.0. The samples were transferred into a 1 mm path length quartz cuvette and analysed on Jasco J-810 DC spectrometer from 198-260nm. Spectra were acquired in triplicate and averaged after subtraction of the buffer background.

### Methylation of *Mtb* UgpB

Purified *Mtb* UgpB was diluted to 1 mg/mL into buffer C and reductively methylated as described previously^41^. Briefly, dimethylborane amine (DMAB) and formaldehyde were added to *Mtb* UgpB and the mixture was left shaking at 100 rpm at 4 °C for two hours. This step was repeated two additional times. DMAB was then added to *Mtb* UgpB for a final incubation step (1 hour, 4 °C, shaking at 100 rpm) followed by the addition of Tris-HCl (final concentration 100 mM, pH 7.0) to remove any excess unreacted DMAB reagent and the sample was then dialysed at 4 °C for 16 hours against buffer C. *Mtb* UgpB was concentrated by ultrafiltration (10 kDa cut-off, Amicon Ultra) to 7 mg/mL.

### Crystallisation and structure determination

For co-crystallisation experiments methylated *Mtb* UgpB was incubated with 10mM glycerol-3-phosphocholine (GPC) and incubated at 4 °C for 30 min before crystallization. Crystals of *Mtb* UgpB in complex with GPC were grown by vapor diffusion in 96-well plates (Swiss-Ci) using a Mosquito liquid handling system (TTP LabTech) by mixing 1:1 volumes (150 nL) of concentrated methylated *Mtb* UgpB (7 mg/mL) with reservoir solution. *Mtb* UgpB crystals typically grew within three days at 22 °C in 0.2 M MgCl2, 0.1 M Tris pH 8.5, 20% w/v PEG 8,000. The *Mtb* UgpB crystals were cryoprotected with 20% glycerol and flash frozen in liquid nitrogen prior to data collection.

The X-ray diffraction data for the ligand bound *Mtb* UgpB crystals were collected at the I04 beamline of Diamond Light Source. The diffraction data were indexed, integrated and scaled with XDS ^42^ through the XIA2 pipeline and the CCP4 suite of programmes ^43^. Initial phases were determined by molecular replacement using PHASER ^44^ and the separate domains (Domain I residues 34-153 and 305-378/ Domain II residues 154304 and 379-424) of the *apo-Mtb* UgpB structure as two ensembles as a search model (PDB 4MFI) specifying to search for 4 copies in the asymmetric unit. Autobuild ^45^ was initially used for model building followed by iterative cycles of alternating manual rebuilding in COOT ^46^ and reciprocal space crystallographic refinement with PHENIX-REFINE ^47^ assigning each domain as a separate TLS group. The coordinates for the glycerol-3-phosphocholine ligand were downloaded from the PDB and fitted into unoccupied electron density in all four chains of the asymmetric unit. The restraints for use in refinement were calculated using REEL ^48^. Magnesium ions and glycerol molecules were also fitted into the unoccupied electron density as well as waters. Methylated lysine (MLZ) was fitted at position 161 in each chain.

The model of the ligand-bound structure comprises residues 36-428 in all chains (A-D), with an additional 1 residue in chains B and D and 2 residues in chain C. There is one disordered region between residues 355-366 in chains C and D and these residues were not modelled. No Ramachandran outliers were identified and structure validations were done by MolProbity ^49^. Figures were prepared using Pymol (The PyMOL Molecular Graphics System, Version 2.0 Schrödinger, LLC), except for those showing electron density which were prepared using CCP4mg ^50^.

### DynDom Analysis

DynDom (http://fizz.cmp.uea.ac.uk/dyndom/)^19^ was used to determine dynamic domain and hinge regions comparing the refined ligand-bound structure and the previously solved apo structure (PDB 4mfi). Default parameters were used for the analysis: a window length of 5, minimum ratio of inter-domain to intradomain displacement of 1.0 and minimum domain size of 20 residues.

### ^1^H STD NMR experiments

All the STD NMR experiments were performed in PBS D2O buffer, pH 7.5. For the complex *Mtb* UgpB/GPC the protein concentration was 68 μM while the ligand concentration was 5 mM. STD NMR spectra were acquired on a Bruker Avance 500.13 MHz at 298 K. The on- and off-resonance spectra were acquired using a train of 50 ms Gaussian selective saturation pulses using a variable saturation time from 0.5 s to 4 s, and a relaxation delay (D1) of 4 seconds. The water signal was suppressed using the watergate technique ^51^ while the residual protein resonances were filtered using a T_1ρ_-filter of 50 ms. All the spectra were acquired with a spectral width of 5 kHz and 32K data points using 128 scans. The on-resonance spectra were acquired by saturating at 0.77 or 6.78 ppm while the off-resonance spectra were acquired by saturating at 40 ppm. Instead, for the *Mtb* UgpB/GPIP4 complex, the protein concentration was 35 μM while the ligand concentration was 2.5 mM. STD NMR spectra were acquired on a Bruker Avance 800.23 MHz at 278 K. The on- and off-resonance spectra were acquired using a train of 50 ms Gaussian selective saturation pulses using a variable saturation time from 0.5 s to 4 s and a relaxation delay (D1) of 5 seconds. The water signal was suppressed by using the excitation sculpting technique ^52^ while the residual protein resonances were filtered using a T_1ρ_-filter of 24 ms. All the spectra were acquired with a spectral width of 12.82 kHz and 32K data points using 64 scans. The on-resonance spectra were acquired by saturating at 0.7 or 6.67 ppm while the off-resonance spectra were acquired by saturating at 40 ppm. To get accurate structural information from the STD NMR data and in order to minimize the T1 relaxation bias, the STD build up curves were fitted to the equation STD(t_sat_) = STD_max_*(1-exp(-k_sat_*t_sat_) calculating the initial growth rate STD0 factor as STD_max_*k_sat_ = STD0 and then normalizing all of them to the highest value ^22^.

### CORCEMA-ST calculations

The CORCEMA-ST software was used to calculate the theoretical STD intensities from the crystallographic structure of the *Mtb* UgpB/GPC complex. The parameters used for the calculations were: saturation frequency range 0-1.1 ppm; protein correlation time 45 ns; K_d_ 0.005 mM; order parameter 0.85; ligand correlation time 0.3 ns; ρ-leak 0.35 s; τ_m_ 10 ps; cutoff 8 Å; [L]0 5 mM; [E]0 68μM; field 500 MHz. While for the *Mtb* UgpB/GPIP4 complex model obtained from docking calculations, the theoretical STD intensities were calculated using the following parameters: saturation frequency range 0-0.9 ppm; protein correlation time 45 ns; K_d_ 1 mM; order parameter 0.85; ligand correlation time 0.3 ns; ρ-leak 0.1 s; τm 10 ps; cutoff 8 Å; [L]0 5 mM; [E]0 68 μM; field 800 MHz. The calculations were repeated in order to have the best fitting possible between the calculated and the experimental ^1^H STD NMR intensities. For CH2 protons showing the same chemical shift an averaged calculated ^1^H STD NMR intensity was assumed. NOE factor ^25^ was used to evaluate the best fit to the experimental data.

### Autodock Vina Docking calculations

Autodock tools ^53^ was used to prepare for docking both the ligand GPIP4 and the *Mtb* UgpB protein. The calculations were performed by positioning a grid of 20 × 24 × 22 Å in the center of the binding site of *Mtb* UgpB, which was maintained rigid while the ligand was considered flexible. The calculations were performed using Autodock Vina ^27^

### DEEP-STD NMR

DEEP-STD factors were obtained as previously described ^24^. Briefly, frequencies derived from shiftx2 ^54^ for aliphatic and aromatic residues present in the binding site of *Mtb* UgpB were used for the position of the saturating selective pulse. The STD obtained using a saturation of 0.5 seconds on aliphatic or aromatic regions (0.7 or 6.67 ppm, respectively) were then used to calculate the DEEP-STD factors.

### Affinity studies with Microscale thermophoresis (MST)

*Mtb*-UgpB protein was labelled using Monolith His-tag labelling kit RED-Tris-NTA, PBS (PBS supplemented with 0.05 % Tween 20 and a constant concentration of UgpB (50 nM) was used. The compounds were prepared in PBS in the concentration range 0-0.5 M. The samples were loaded into the MonoLith NT.115 standard treated capillaries and incubated for 10 min before analysis using the Monolith NT.115 instrument (NanoTemper Technologies) at 21°C using medium laser power and 40 % LED power. The binding affinities were calculated using a single-site binding model with GraphPad Prism software (version 7.0). All experiments were carried out in triplicate.

### Thermal shift assay

The transition unfolding temperature *T_m_* of the *Mtb* UgpB protein (22 μM) was determined in the presence or the absence of ligands. The screen used a single ligand concentration of 100 mM. Reactions were performed in a total volume of 20 μL using Rotor-Gene Q Detection System (Qiagen), setting the excitation wavelength to 470 nm and detecting emission at 557 nm of the SYPRO Orange protein gel stain, 15 × final concentration (Invitrogen). The cycle used was a melt ramp from 30 to 95°C, increasing temperature in 1°C steps and time intervals of 5 s. Fluorescence intensity was plotted as a function of temperature. The *T_m_* was determined using the Rotor-Gene Q software and the Analysis Melt functionality. All experiments were performed in triplicate.

### Chemoenzymatic synthesis of glycerophosphoethanolamine (GPE) and glycerophosphoserine (GPS)

Phospholipase A1 from *Aspergillus oryzae* was dialysed into PBS overnight at 4°C prior to use. The enzymatic reaction contained either 50 mg of 1,2-Dipalmitoyl-sn-glycero-3-phosphoethanolamine or 25 mg of 1,2-Diacyl-sn-glycero-3-phospho-L-serine in an organic-aqueous media (800 μL hexane, 138 μL H2O: 5.8:1 ratio) and heated at 50°C for 10 mins prior to the addition of Phospholipase A1 from *Aspergillus oryzae* (1 μL Phospholipase A1 per 1 mg of phospholipid). The reaction mixtures were heated at 50°C and stirred at 300 rpm for 48 hours. The solvent was removed *in vacuo* and the reaction mixture redissolved in water (5 mL) and extracted with chloroform (3 × 25 mL). The aqueous phase was separated and the phospholipase A1 enzyme removed using a centrifugal filter unit (Amicon, 10 kDa molecular weight cut off). The collected filtrate was concentrated *in vacuo* to give the products as a colourless oil (9.4 mg GPE) or yellow oil (3 mg GPS). GPE: ‘H NMR (400MHz, D2O) δ_ppm_ 3.99 – 4.08 (2H, m, POC<u>H</u>2CH2N), 3.75 – 3.91 (3H, m, POC<u>H</u>2C<u>H</u>CH2), 3.49 – 3.63 (2H, m, POCH2CHC<u>H</u>2), 3.20 (2H, t, *J* = 5.0 Hz, POCH2C<u>H</u>2N). ^13^C NMR (100MHz, D2O) δ_ppm_ 70.7 (<u>C</u>H), 66.5 (O<u>C</u>H2), 62.0 (O<u>C</u>H2), 61.8 (O<u>C</u>H2), 40.0 (N<u>C</u>H2). ^31^P NMR (161 MHz, D2O) δ_ppm_ 0.42. GPS: ‘H NMR (400MHz, D2O) δ_ppm_ 4.17 – 4.26 (2H, m, POC<u>H</u>2CHN), 4.01 – 4.08 (1H, m, POCH2C<u>H</u>N), 3.71 – 3.93 (3H, m, 3H, m, POC<u>H</u>2C<u>H</u>CH2), 3.48-3.65 (2H, m, POCH2CHC<u>H</u>2). ^13^C NMR (100MHz, D2O) δ_ppm_ 178.5 (<u>C</u>=O), 70.6 (CH2<u>C</u>H(OH)CH2), 66.5 (O<u>C</u>H2), 63.7 (PO<u>C</u>H2CHN), 62.0 (O<u>C</u>H2), 54.4 (POCH2<u>C</u>HN). ^31^P NMR (161 MHz, D2O) δ_ppm_ 0.08.

### Deposition of Coordinates and Structure Factors

Coordinates and structure factors for *Mtb* UgpB have been deposited in the Protein Data Bank under accession code XXXX.

## Acknowledgements

We would like to thank Mohd Syed Ahanger for technical assistance. We thank Professor William R. Jacobs (Albert Einstein College of Medicine, USA) for providing expression vector pYUB1062 and the *M. smegmatis* mc^2^4517 expression system. We acknowledge the contribution of the WPH Proteomics Facility research technology platform in the School of Life Sciences, University of Warwick. We thank Diamond Light Source for access to synchrotron beamlines and their staff for support during experiments. Equipment was supported through the Warwick Integrative Synthetic Biology (WISB) research technology platform (Grant reference BB/M017982/1). This work was supported by a Sir Henry Dale Fellowship to EF jointly funded by the Wellcome Trust and Royal Society (Grant number 104193/Z/14/Z), a research grant from the Royal Society (Grant number RG120405), the MRC for a studentship to JF (grant number MR/J003964/1) and the EPSRC for funding JH as an Integrate Early Career fellow (grant number EP/M027503/1). JA and RN acknowledge support by the Biotechnology and Biological Sciences Research Council (BBSRC) through a New Investigator grant awarded to JA (BB/P010660/1). We are grateful for the use of the University of East Anglia (UEA) Faculty of Science NMR facility.

