## Supporting information for "Structural basis of glycerophosphodiester recognition by the *Mycobacterium tuberculosis* substrate-binding protein UgpB"

**Fig. S1. SDS-PAGE analysis of the purification of *Mtb* UgpB from *M. smegmatis*.** A) Elution of $His_6$-tagged *Mtb* UgpB from a $Co^{2+}$ IMAC-column. M = molecular weight marker in kDa, IS = insoluble fraction, S = soluble lysate, FT = flow through, numbers 5 – 1000 refer to the imidazole concentration in the elution buffer (units of mM). B) QHP anion exchange chromatography of *Mtb* UgpB following the $Co^{2+}$ IMAC step. L = protein after dialysis, FT1 = first flow through, FT2 = second flow through, numbers 150 – 1000 refer to NaCl concentration in the elution buffer (units of mM). C) Size exclusion chromatography of *Mtb* UgpB following anion exchange chromatography with the volumes shown as corresponding to D. D) Size exclusion trace of *Mtb* UgpB. See Materials and Methods for buffer compositions.

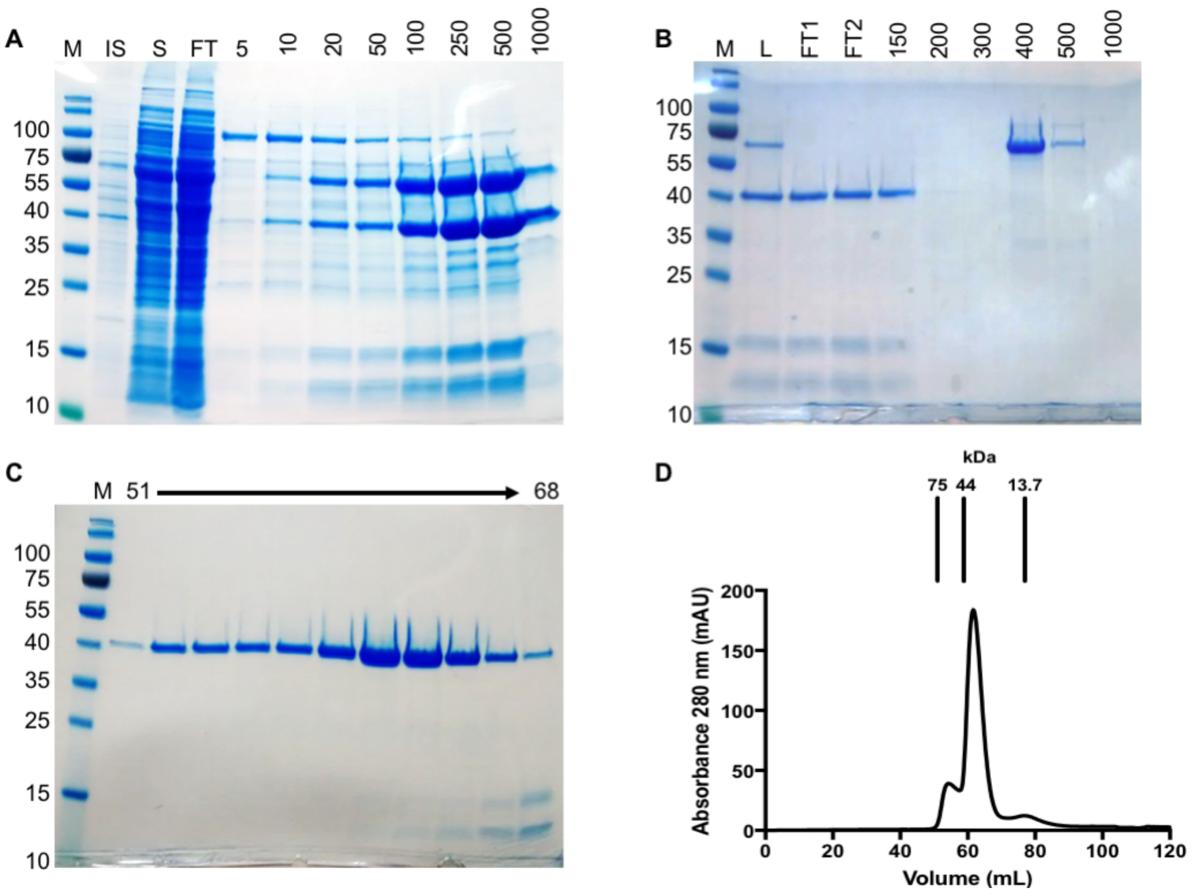

**Fig S2: GPC binding**

**A) Electron density for the GPC substrate.** Electron density map contoured at 0.38electrons/Å$^3$. Carbon atoms are shown in green, oxygen atoms are shown in red, nitrogen atoms in blue and phosphate atom in purple. The figure was prepared using CCP4mg. The .mtz file was loaded directly with the default settings and clipped to select for the GPC atoms.

**B) Alignment of the glycerophosphocholine ligand from each *Mtb* UgpB subunit.** Superposition of GPC from each *Mtb* UgpB subunit in the asymmetric unit. The GPC ligand is shown with green carbon atoms (subunit A), light blue carbon atoms (subunit B), wheat carbon atoms (subunit C) and grey carbon atoms (subunit D). In all subunits oxygen atoms are coloured red and the phosphorous atom is coloured orange.

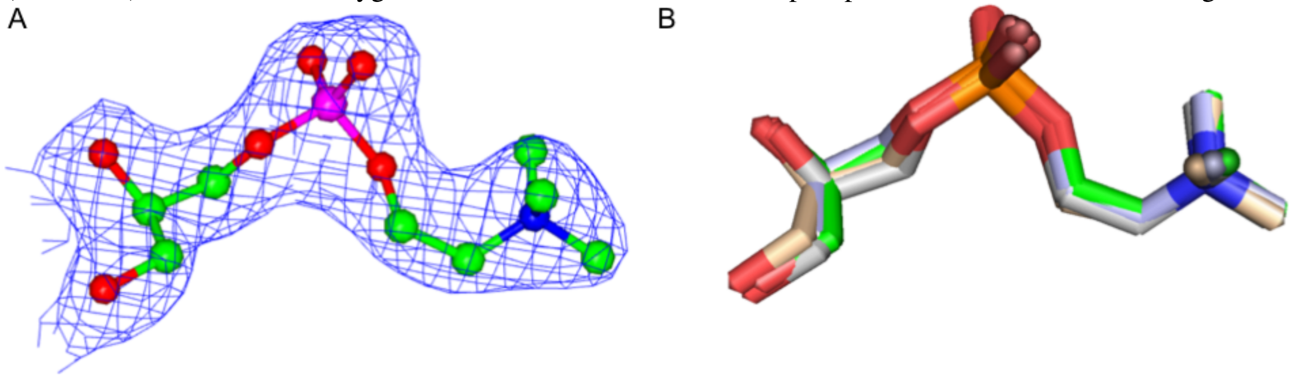

**Fig. S3. Sequence alignment of UgpB from *Mycobacterium tuberculosis* with the UgpB homologue from *Escherichia coli.*** The sequence alignment was generated using Clustal Omega (https://www.ebi.ac.uk/Tools/msa/clustalo/) and ESPript version 3. Identical residues are indicated by a red background and conserved residues by red characters. The secondary structure elements of *Mtb* UgpB are shown above the sequences and the secondary structure elements of *E. coli* UgpB (4AQ4) are shown below the sequences.

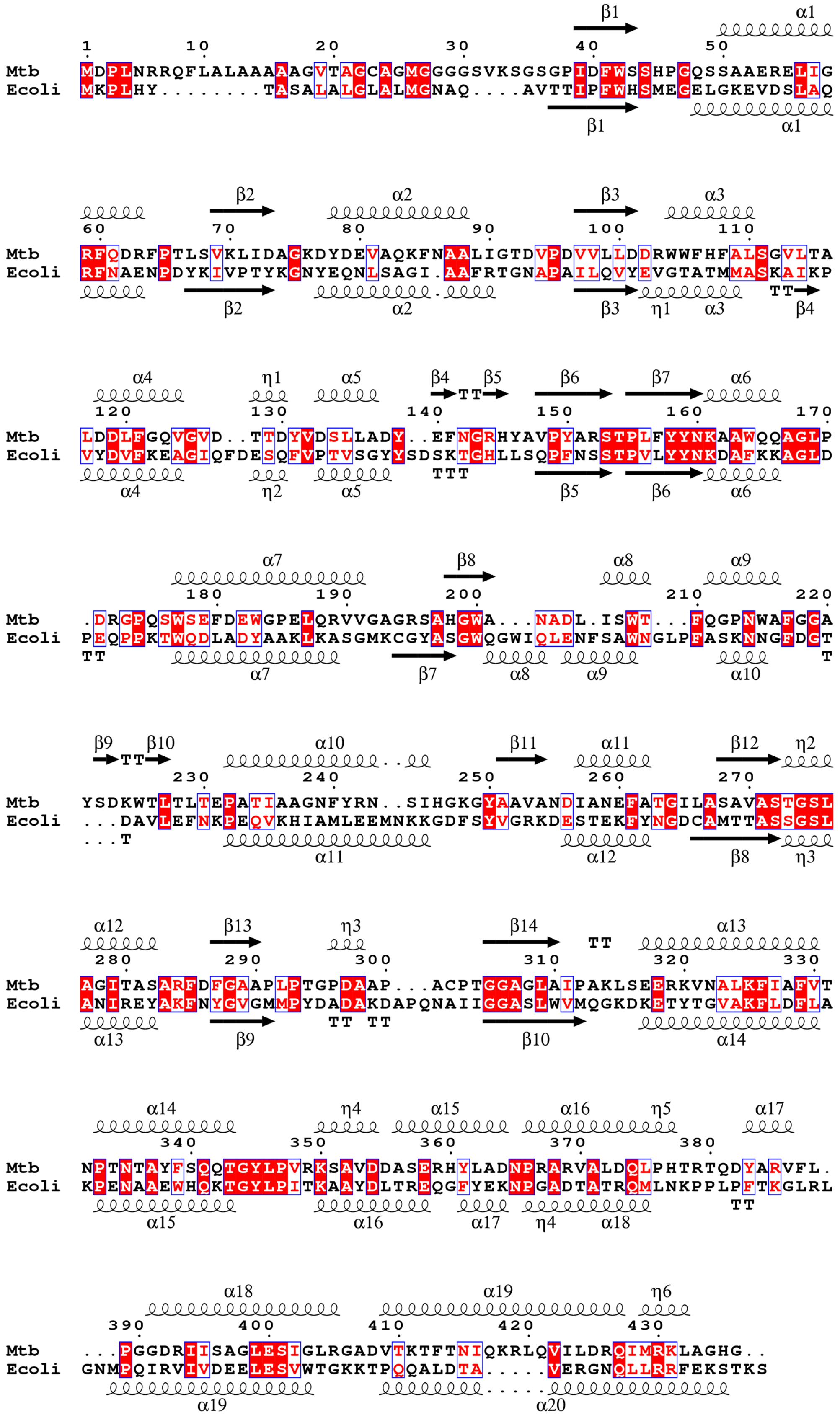

**Fig S4: Location of additional glycerol moiety in the *Mtb* UgpB binding pocket.** A) Surface representation of *Mtb* UgpB. The GPC ligand is represented by yellow spheres and a solvent glycerol moiety as green spheres and the. B) Close-up of the *Mtb* UgpB binding pocket with the GPC ligand and glycerol moiety shown in stick representation.

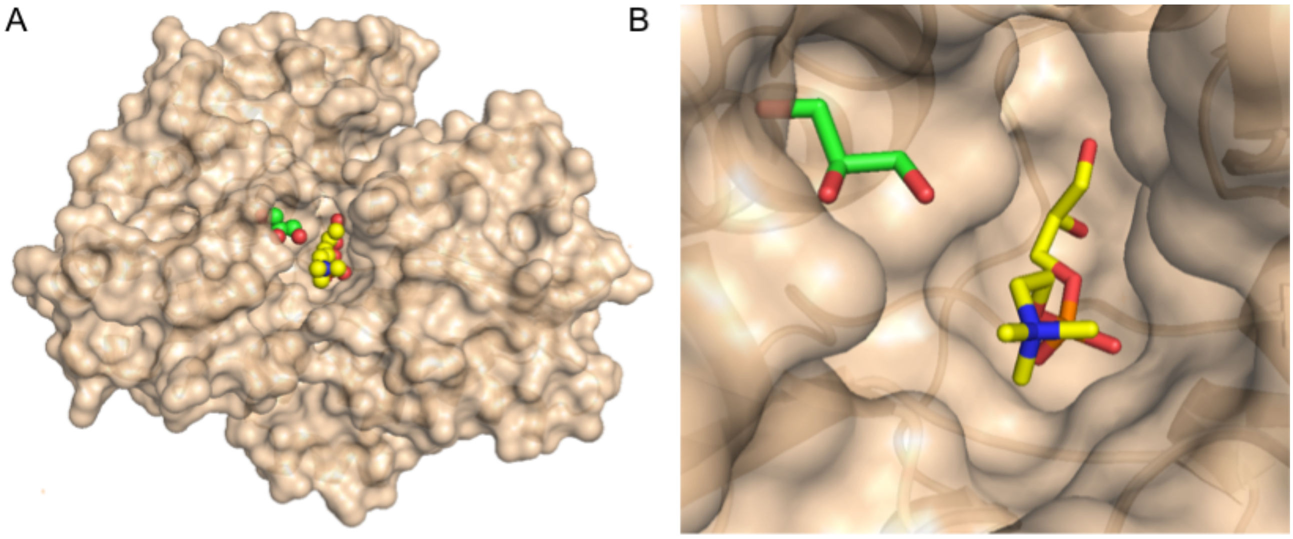

**Fig S5. Thermal shift assay probing a panel of potential *Mtb* UgpB ligands.** Bar graphs illustrating shifts of $\Delta T_m$ for the series of potential ligands. Thirty seven different ligands were probed for binding at a final concentration of 100mM. Data are shown from three independent repeats represented as mean ± SD.

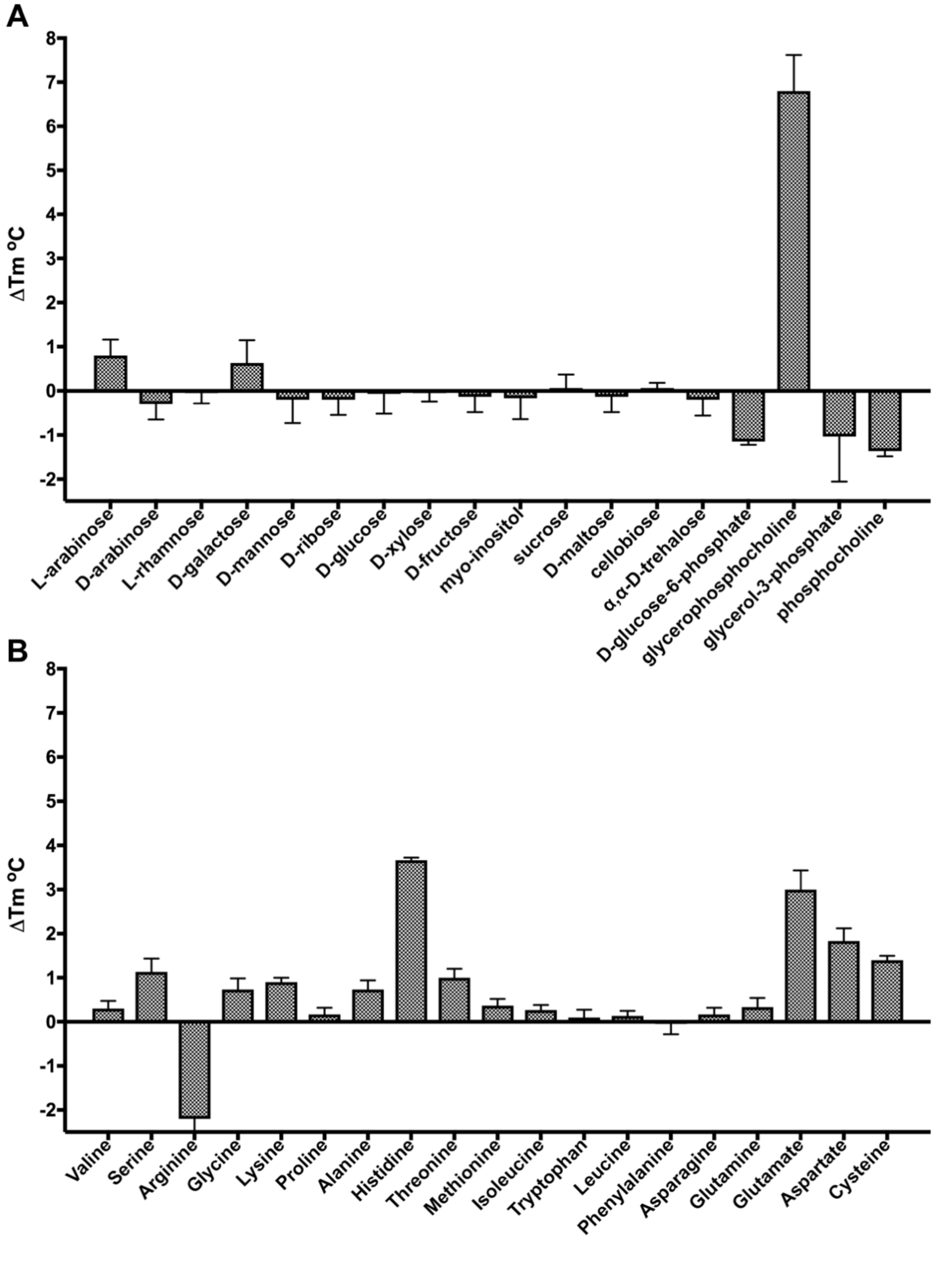

**Fig S6. CD spectra of *Mtb* UgpB and site directed mutant proteins.** CD spectra of *Mtb* UgpB (red)), *Mtb* UgpB Tyr78Ala (green), *Mtb* UgpB Asp102Ala (cyan), *Mtb* UgpB Ser153Ala (purple), *Mtb* UgpB Leu205Ala (magenta), *Mtb* UgpB Trp208Ala (brown), *Mtb* UgpB Ser272Ala (orange), Tyr345Ala (blue), *Mtb* UgpB Arg385Ala (yellow).

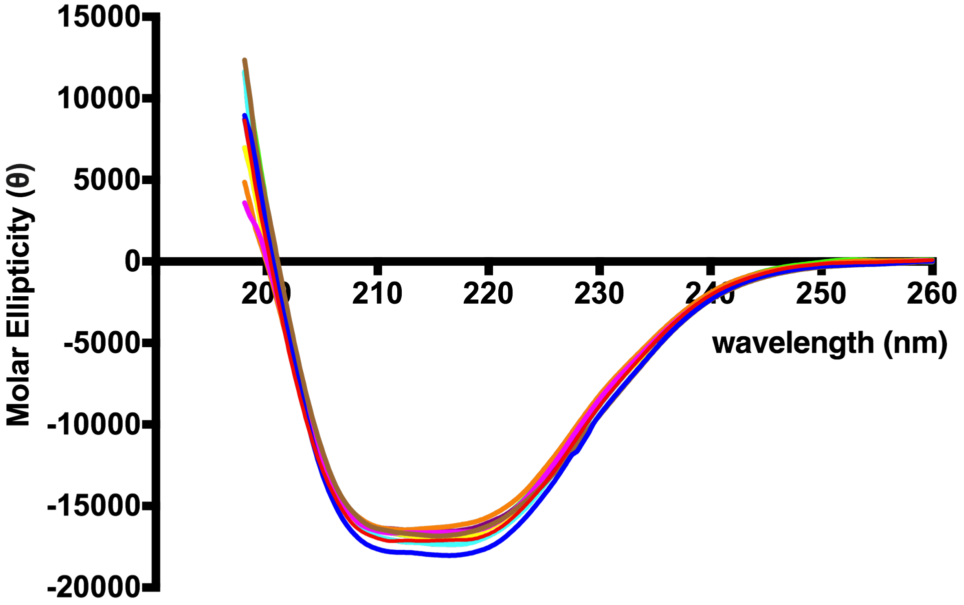

**Table S1. DynDom analysis of *Mtb* UgpB (pdb 4MFI) and *Mtb* UgpB in complex with GPC**

| Backbone RMSD (Å) | Bending region | Rotation angle | Translation (Å) | Closure (%) |
|---|---|---|---|---|
| 0.47 (Domain I) | 152-153 | 21.8º | 0.8 | 98.7 |
| 0.54 (Domain II) | 304-306 | | | |
| | 362-372 | | | |

Domain I comprises residues 38-152 and 306-365 and Domain II comprises residues 153-305 and 366-426

**Table S2. Sequence of primers for cloning and site-directed mutagenesis**

Restriction recognition sites are in italics. The codon encoding the amino acid mutation is indicated in bold type.

| Name | Use | Sequence (5’-3’) |
|---|---|---|
| UgpB_T_pYUB_5 | Clone truncated *Mtb* UgpB pYUB1062 | aaaaaa*catatg*ggttccggcccaatcgacttctgg |
| UgpB_T_pYUB_3 | Clone truncated *Mtb* UgpB pYUB1062 | aaaaaa*aagctt*gccatgccccgccagcttccg |
| Tyr78Ala_F | Mutate *Mtb* UgpB residue Tyr78Ala | ggcaaggac**gcc**gacgaggtg |
| Tyr78Ala_R | Mutate *Mtb* UgpB residue Tyr78Ala | cacctcgtc**ggc**gtccttgcc |
| Asp102Ala_F | Mutate *Mtb* UgpB residue Asp102Ala | cgttttgctcgac**gcc**cgatggtggttcc |
| Asp102Ala_R | Mutate *Mtb* UgpB residue Asp102Ala | ggaaccaccatcg**ggc**gtcgagcaaaacg |
| Ser153Ala_F | Mutate *Mtb* UgpB residue Ser153Ala | ccgtatgctcgc**gcg**acgccgctgttc |
| Ser153Ala_R | Mutate *Mtb* UgpB residue Ser153Ala | gaacagcggcgt**cgc**gcgagcatacgg |
| Leu205Ala_F | Mutate *Mtb* UgpB residue Leu205Ala | gctaacgccgac**gcc**atctcgtggacg |
| Leu205Ala_R | Mutate *Mtb* UgpB residue Leu205Ala | cgtccacgagat**ggc**gtcggcgttagc |
| Trp208Ala_F | Mutate *Mtb* UgpB residue Trp208Ala | ccgacctcatctc**ggc**gacgtttcagggacc |
| Trp208Ala_R | Mutate *Mtb* UgpB residue Trp208Ala | ggtccctgaaacgtc**gcc**gagatgaggtcgg |
| Ser272Ala_F | Mutate *Mtb* UgpB residue Ser272Ala | gccgtggca**gcc**accggctcg |
| Ser272Ala_R | Mutate *Mtb* UgpB residue Ser272Ala | cgagccggt**ggc**tgccacggc |
| Tyr345Ala_F | Mutate *Mtb* UgpB residue Tyr345Ala | cagcaaaccggc**gct**ctgccggtgcgcaag |
| Tyr345Ala_R | Mutate *Mtb* UgpB residue Tyr345Ala | cttgcgcaccggcag**agc**gccggtttgctg |
| Arg385Ala_F | Mutate *Mtb* UgpB residue Arg385Ala | cacaagactacgca**gcg**gttttcctgcc |
| Arg385Ala_R | Mutate *Mtb* UgpB residue Arg385Ala | ggcaggaaaac**cgc**tgcgtagtcttgtg |
